# Social stress, cortisol awakening response and sex: Impact on hippocampus and amygdala volume

**DOI:** 10.1101/2025.03.20.644300

**Authors:** Rui Wang, Hannes Noack, Leandra Kuhn, Vanessa Nieratschker, Ute Habel, Birgit Derntl, Lydia Kogler

**Affiliations:** Department of Psychiatry and Psychotherapy, Women’s Mental Health & Brain Function, Tübingen Center for Mental Health (TüCMH), Medical Faculty, University of Tübingen, Tübingen, Germany; Department of Psychiatry, Psychotherapy and Psychosomatics, RWTH Aachen University, Aachen, Germany; Department of Psychiatry and Psychotherapy, Molecular Psychiatry, Tübingen Center for Mental Health (TüCMH), Medical Faculty, University of Tübingen, Tübingen, Germany; Institute of Neuroscience and Medicine (INM-10), Research Centre Jülich, Jülich, Germany; German Center for Mental Health (DZPG), partner site Tübingen, 72076 Tübingen, Germany; LEAD Graduate School and Research Network, University of Tübingen, 72074 Tübingen, Germany

**Author notes:** Corresponding author at: Department of Psychiatry and Psychotherapy, University of Tübingen, Calwerstraße 14, 72076 Tübingen, Germany. E-mail address (RW); (LK).

**Keywords:** hippocampus, amygdala, gray matter volume, sex difference, cortisol awakening response, social stress

## Abstract

**Objective:** Volumes of the hippocampus and amygdala, both major hubs for neural stress regulation amongst others, are influenced by social stressors, cortisol awakening response (CAR) and sex, i.e. psychological and physiological stress factors. Importantly, the interplay of these different factors in affecting the morphology of both brain regions remains unclear. This study aimed to elucidate the intricate influence of these factors on grey matter volumes (GMV) of the hippocampus and amygdala.

**Methods:** We analyzed associations between structural brain data, self-reported chronic social stress (including subscales on social tension, social overload, lack of social recognition and social isolation) and CAR of 83 healthy participants (40 females) with multiple regression analyses.

**Results:** In males, but not females, higher social tension predicted lower bilateral hippocampus GMV. In individuals not showing the typical CAR, higher social tension predicted reduced bilateral hippocampus GMV. Amygdala GMV was also related to social stress and CAR, with social overload associated with reduced amygdala GMV in individuals not showing the typical CAR, while the opposite pattern emerged in those with an atypical CAR.

**Conclusions:** Chronic social stress is associated with hippocampus and amygdala volumes, interacting with CAR and sex. The brain morphology in males and in individuals with atypical CAR seems to be more vulnerable to chronic social stress. Our findings enhance our understanding of how brain architecture, social stress, CAR patterns and sex shape individual stress responses.

## 1. Introduction

Stress reactivity involves a distributed neural circuitry, with key brain regions including the hippocampus, amygdala, and prefrontal cortex playing central roles in adaptive responses to stressful situations (McEwen et al., 2016; McEwen and Gianaros, 2010). However, when stress is prolonged, such as in chronic social stress, these regions, particularly the hippocampus and amygdala, are adversely affected (Fuchs and Flügge, 2003; McEwen et al., 2016). Structural reductions in these areas have been observed in rats exposed to repeated social defeats (Patel et al., 2018), as well as in humans subjected to early-life stress (Hanson et al., 2015) and stress in the past months (Piccolo and Noble, 2018; Sublette et al., 2016; Zimmerman et al., 2016). Furthermore, reductions in their volumes are linked to stress-associated disorders, such as depression (Nolan et al., 2020) and post-traumatic stress disorder (Del Casale et al., 2022). These findings emphasize the importance of the hippocampus and amygdala in stress responses and indicate their vulnerability to chronic stress.

Both brain regions are significantly influenced by glucocorticoids (primarily cortisol in humans), due to their high density of gluco- and mineralocorticoid receptors (De Kloet et al., 1998; Wang et al., 2014). Cortisol regulates the stress response by binding to these receptors, a process closely linked to the hypothalamus-pituitary-adrenal (HPA) axis, which controls cortisol secretion through a cascade of responses (Herman et al., 2012). In healthy individuals, cortisol exhibits a distinct circadian rhythm, characterized by a significant increase in cortisol levels around 30-45 minutes after awakening in the morning, the so-called cortisol awakening response (CAR), followed by a subsequent decline over the day (Pruessner et al., 1997). Chronic stress is known to disrupt this pattern, leading to a reduced increase in cortisol after awakening (Duan et al., 2013), possibly due to prolonged activation of the HPA axis, which impairs the ability to respond adaptively after awakening. This atypical CAR pattern is thought to reflect impaired neuroendocrine function and dysregulated circadian rhythm (Stalder et al., 2016), which is further related to brain atrophy and reduced volumes of the hippocampus (Pruessner et al., 2017).

Research reports sex differences in reaction to social stress. Females show greater response to social rejection (Stroud et al., 2002) and are more susceptible to exhaustion from social media overload compared to males (Wu and Zheng, 2023). Furthermore, a review by Gray et al. (2017) assessing the associations between glucocorticoids and steroid hormones such as estradiol or testosterone with stress-induced brain changes, further emphasizes the importance of considering sex. Evidence from animal studies demonstrates consistent results: In males, chronic stress, induced by chronic restraint task, causes dendritic atrophy in the hippocampus and the amygdala, whereas females do not show the same pattern (Galea et al., 1997; Lin et al., 2009), unless they are ovariectomized (i.e., significantly decreased estradiol levels) (McLaughlin et al., 2010, 2005). This suggests that ovarian hormones may attenuate the effects of chronic stress on hippocampus dendritic atrophy (Conrad et al., 2012; McLaughlin et al., 2009). In humans, sex-dependent associations between stress-induced cortisol levels and striatal-limbic structures were reported, with positive associations observed in males and negative ones in females (Henze et al., 2023). Additionally, females were observed to have greater functional connectivity within the amygdala network in response to social exclusion compared to males (Bürger et al., 2023); and males showed more pronounced neural activation in the inferior frontal gyrus and insula (Kuhn et al., 2022). In stress-related mental disorders, sex differences appeared with an atypical, reduced CAR going along with smaller left hippocampus volumes in male patients with first-episode psychosis and smaller bilateral hippocampus volumes in high-risk individuals, whereas this association was not seen in females (Pruessner et al., 2015, 2017). So far, it remains unclear whether similar sex-specific patterns between CAR and volumes of stress-regulating brain areas such as the hippocampus and the amygdala appear in healthy individuals as well.

Taken together, the interplay between chronic social stress and CAR on hippocampus and amygdala volumes, as well as the effect of sex on the relationship between chronic social stress and these brain structures remain unclear. The current study aims to explore interactions between chronic social stress, CAR, and sex affecting grey matter volume (GMV) of the hippocampus and the amygdala. More specifically, we want to assess the impact of the following variables on GMV of the hippocampus and amygdala: 1) the interaction between chronic social stress and CAR; 2) the interaction of chronic social stress and sex; and 3) the interaction of sex and CAR. Our hypotheses are as follows: 1) individuals with lower CAR experiencing high chronic social stress will show reduced GMV of the hippocampus and the amygdala, potentially due to their deficient HPA axis regulation (Stalder et al., 2016); 2) in males, but not in females, chronic social stress has a significant negative impact on GMV of the hippocampus and amygdala (Galea et al., 1997; Lin et al., 2009); and 3) in males, but not in females, a lower CAR is associated with reduced hippocampus volumes (Pruessner et al., 2017, 2015).

## 2. Methods

### 2.1. Sample

Participants were part of a multi-centric study (Kuhn et al., 2022, 2021), recruited through public advertisements at the universities of Tuebingen and RWTH Aachen University, Germany and were between 18 and 35 years of age. Eligibility for the study was evaluated through a semi structured telephone interview and the screening version of the German Structured Clinical Interview for Diagnostic and Statistical Manual of Mental Disorders (SCID, Wittchen et al., 1997). Inclusion criteria were right-handedness, nonsmokers, normal weight (17 < BMI < 30), a regular day-night cycle (no shift working, jet lag). Exclusion criteria included any self-reported history of neurological or mental disorders, pregnancy, as well as common requirements for MRI scanning, such as no metal parts in or on the body, tattoos, pacemakers or similar, etc. Additionally, only females using oral contraceptives were included during their pill-intake phase to minimize the impact of hormonal fluctuations on cortisol reactivity (Kirschbaum et al., 1999). Participants completed the State-Trait Anxiety Inventory (STAI, Spielberger et al., 1980) and the Beck Depression Inventory-II (BDI, Beck, 1996; Jackson-Koku, 2016) to assess their anxiety and depression levels.

Of note, the term “sex” used throughout this paper refers to the biological sex assigned at birth, as reported by participants, and is categorized as female or male in accordance with the Sex and Gender Equity in Research (SAGER) guidelines (Heidari et al., 2016).

The study protocol was approved by the ethics committees of the medical faculties at the RWTH Aachen University (EK 212/16) and the University of Tuebingen (409/2015BO2), and was conducted in accordance with the declaration of Helsinki (World Medical Association, 2013).

### 2.2. Cortisol collection

Saliva samples were collected on a normal working-day with a typical daily routine using Cortisol Salivettes (Sarstedt, Nürnbrecht, Germany) immediately after awakening, 15, 30, 45 and 60 min post-awakening. Participants received detailed instructions during the screening appointment a few days before the MR session and returned the samples when they came for the MR session. Participants placed a synthetic fiber swab in their mouth for one minute to absorb saliva and were told not to chew the swab. Participants recorded the exact time of each sample collection to ensure adherence to the timing protocol. They were instructed not to brush their teeth, drink, eat or smoke prior to the sampling procedures, to abstain from alcohol and nicotine consumption and excessive exercise on the day before sample collection, and to maintain their usual waking times and daily routines on the sampling day. After collection, participants stored the samples in a fridge and brought them to the laboratory in the afternoon of the same day, when the MR-scanning and the assessment of chronic social stress took place. The samples were kept in the refrigerator or freezer until they were sent for analysis to the Institute of Pharmaceutical Sciences, University of Tuebingen.

Salivary cortisol was measured using a liquid chromatography (LC)-mass spectrometry assay. Samples preparation involved: thawing at 4 °C, protein precipitation, solid-phase extraction using Oasis PRIME hydrophilic-lipophilic balance material on a 96-well plate, drying under nitrogen, and reconstituting in 50 μL of a methanol-water solution (30:70; v/v) in a sealed 96-well collection plate. Chromatographic separation was performed using a MicroLC 200 Plus instrument (Sciex, Framingham, MA, USA), with analyte detection carried out on a QTRAP 4500 triple-quadrupole mass-spectrometer (Sciex) in negative ionization mode. For accurate quantification, a surrogate calibrant method using Cortisol-d4 in a true saliva matrix was employed. The method demonstrated a linear range for cortisol quantification from 0.062 to 75.5 ng/mL and was validated in accordance with FDA guidelines.

### 2.3. Chronic social stress

To assess chronic social stress levels, we investigated subscales of the Trier Inventory for the Assessment of Chronic Stress (TICS, Schulz et al., 2004; Schulz and Schlotz, 1999), assessing stress resulting from high social demands or a lack of social need satisfaction appearing within the previous 3-months: “social overload” (the feeling of being overwhelmed due to the number and intensity of social interactions and responsibilities); “lack of social recognition” (not receiving sufficient appreciation or recognition from others in social situations); “social tension” (conflicts and tense interactions in social networks); and “social isolation” (feelings of loneliness and isolation from others).

### 2.4. MRI data acquisition and pre-processing

For structural magnetic resonance imaging (MRI) data of the brain, participants were scanned with a T1-weighted magnetization prepared rapid acquisition gradient-echo (MPRAGE) sequence. The acquisition parameters were as follows: TR = 5,000 ms, TE = 2.98 ms, flip angle = 9◦, FOV = 256 × 256 mm, 176 slices, voxel size = 1 mm^3^, interleaved, with a distance factor of 50%.

The preprocessing of T1-weighted images was performed by using the Computational Anatomy Toolbox (CAT12, https://neuro-jena.github.io/cat/), the Statistical Parametric Mapping software (SPM12, https://www.fil.ion.ucl.ac.uk/spm/) and MATLAB R2023a (Mathworks, Natick, MA, USA, https://uk.mathworks.com/). The cross-sectional data segmentation tool was run with the default settings provided by CAT12. Each participant’s T1-weighted image underwent spatial normalization and segmentation into grey matter (GM), white matter (WM) and cerebrospinal fluid (CSF). These images were normalized to a standard stereotactic space (MNI template), with a resulting voxel size of 2 mm for the normalized images. After preprocessing, the modulated and normalized GMV images for each participant were smoothed by an 8 mm full width at half maximum (FWHM) isotropic Gaussian kernel.

For region of interest (ROI) analyses, masks were created based on the neuromorphometrics atlas using the Image Calculator tool in SPM12. The location of the ROIs in this study is shown in Figure 1. GMVs were extracted from the left and right hippocampus, and the left and right amygdala by “get_totals script, G. Ridgeway, http://www0.cs.ucl.ac.uk/staff/gridgway/vbm/get_totals.m, date of access: June 27, 2023”, as done in a previous study (Pletzer et al., 2019).

**Figure 1.**
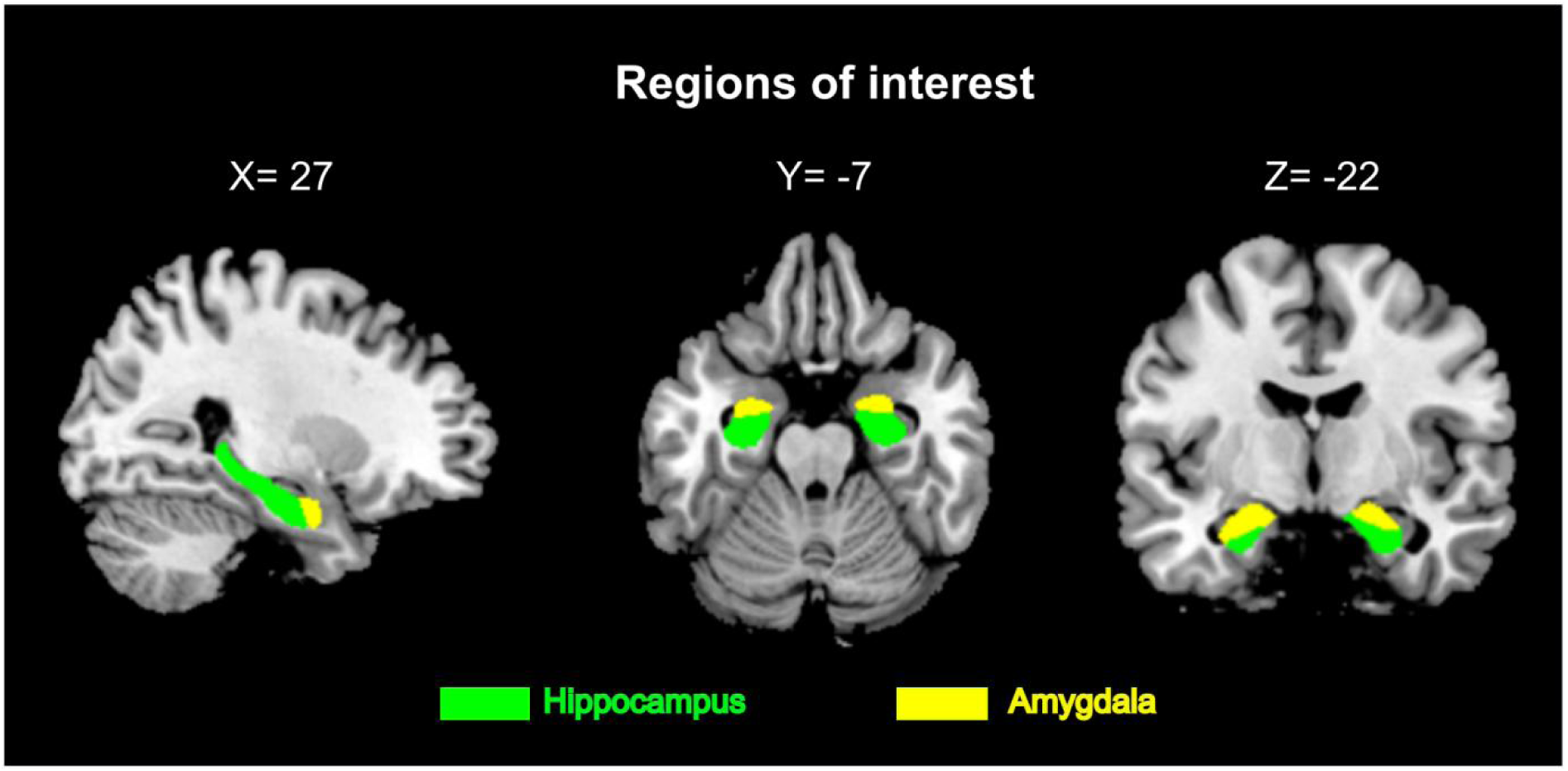
Regions of interest in MNI space at coordinate X = 27, Y = −7, Z = −22.

### 2.5. Statistical analyses

Data were analyzed using IBM SPSS Statistics 29 (https://www.ibm.com/de-de/products/spss-statistics). Based on established criteria, participants were categorized into responders and non-responders according to their cortisol increase at 30 minutes after awakening. Responders demonstrated an increase exceeding 50%, whereas non-responders exhibited an increase below 50% (Clow et al., 2004; Xiong et al., 2021).

Multiple regression analyses were performed to examine whether chronic social stress, CAR and sex are predictive of GMV. Separate analyses were conducted for the left and right hippocampus and amygdala as dependent variables. For each side of each ROI, four different models were applied corresponding to four subscales of chronic social stress. The predictors in the regression model were sex (females/males), CAR (responders/non-responders), chronic social stress level, and their interactions (chronic social stress × sex; chronic social stress × CAR; sex × CAR). Total intracranial volume (TIV) and age were included as covariates of no interest in each model. In cases where significant interactions were identified, partial correlation analyses were performed, controlling for TIV and age, to clarify the associations within each group. Additionally, Fisher’s Z transformation was used to compare differences in correlation coefficients between groups. To reduce multicollinearity, chronic social stress scores were centered. The CAR group and sex were dummy-coded. To control for multiple comparisons in the left and right hemispheres, p-values were adjusted using Bonferroni correction, with a significance threshold set at 0.025.

## 3. Results

### 3.1. Sample characteristics

A total of 83 participants (40 females) were included in the current study based on the data availability within the original database. Of these, 45 participants (22 females) were classified as responders and 38 participants (18 females) as non-responders according to the increasing levels of cortisol after awakening. Consequently, responders showed higher area under the curve with respect to the increase (AUCi) in the CAR compared to non-responders (t(81) = 3.63, p < 0.001), aligned with the classification criteria. No other significant differences were shown between groups. Moreover, results of the chi-square test indicated that there was no significant difference in the distribution of CAR responders and non-responders between females and males (χ² = 0.019, p = 0.99). Additionally, males were older compared to females (t(81) = 2.14, p = 0.035), and as expected, males exhibited a larger TIV than females (t(81) = 8.653, p < 0.001). No further significant sex differences in sample characteristics were observed. Please see Table 1 for details.

**Table 1:** Sample characteristics.

|  | **Overall**<br>**(N = 83)** | **Females**<br>**(N = 40)** | **Males**<br>**(N = 43)** | **p<sup>1</sup>** | **Responders**<br>**(N = 45)** | **Non-responders**<br>**(N = 38)** | **p<sup>2</sup>** |
| --- | --- | --- | --- | --- | --- | --- | --- |
| Age (years) | 23.22 ± 3.11 | 22.48 ± 2.56 | 23.91 ± 3.43 | **0.035** | 22.64 ± 2.98 | 23.89 ± 3.16 | 0.068 |
| BDI-II | 2.99 ± 3.34 | 2.83 ± 3.45 | 3.14 ± 3.26 | 0.671 | 3.04 ± 3.46 | 2.92 ± 3.22 | 0.868 |
| STAI-State | 43.41 ± 7.03 | 44.24 ± 7.50 | 42.66 ± 6.57 | 0.323 | 42.45 ± 7.08 | 44.53 ± 6.89 | 0.195 |
| STAI-Trait | 33.61 ± 7.78 | 33.73 ± 7.26 | 33.50 ± 8.34 | 0.897 | 33.16 ± 7.11 | 34.13 ± 8.56 | 0.576 |
| Cortisol after awakening (ng/mL) |  |  |  |  |  |  |  |
| CAR - AUCg | 475.96 ± 655.17 | 517.64 ± 754.39 | 438.17 ± 556.65 | 0.679 | 450.69 ± 592.72 | 506.70 ± 731.20 | 0.703 |
| CAR - AUCi | 114.99 ± 304.86 | 117.08 ± 243.41 | 113.08 ± 354.43 | 0.367 | 218.32 ± 345.52 | -10.69 ± 183.34 | **<0.001** |
| Awakening time (a.m.) | 7:27 ± 1:11 | 7:16 ± 1:13 | 7:41 ± 1:07 | 0.190 | 7:15 ± 1:05 | 7:43 ± 1:12 | 0.051 |
| Sleep duration (hours) | 7.68 ± 1.22 | 7.87 ± 1.24 | 7.45 ± 1.17 | 0.132 | 7.60 ± 1.13 | 7.57 ± 1.23 | 0.917 |
| Chronic social stress |  |  |  |  |  |  |  |
| Social overload | 7.00 ± 4.77 | 6.63 ± 4.42 | 7.35 ± 5.10 | 0.493 | 6.36 ± 4.29 | 7.76 ± 5.24 | 0.182 |
| Lack of social recognition | 4.01 ± 2.72 | 3.98 ± 2.65 | 4.05 ± 2.81 | 0.905 | 3.67 ± 2.59 | 4.42 ± 2.83 | 0.209 |
| Social tension | 5.25 ± 4.66 | 5.00 ± 4.82 | 5.49 ± 4.54 | 0.636 | 4.89 ± 4.42 | 5.68 ± 4.94 | 0.442 |
| Social isolation | 6.36 ± 4.23 | 5.88 ± 3.76 | 6.81 ± 4.63 | 0.316 | 6.31 ± 4.42 | 6.42 ± 4.06 | 0.907 |
| TIV | 1563.55 ± 158.48 | 1450.36 ± 116.78 | 1668.85 ± 113.22 | **<0.001** | 1569.80 ± 165.93 | 1556.16 ± 151.04 | 0.699 |

Note. Values are presented as mean ± standard deviation. Sleep data were from 56 participants due to missing sleep data for some participants.
<sup>1</sup> indicates p-value for the differences between females vs. males. <sup>2</sup> indicates p-value for the differences between responders vs. non-responders.
BDI = Beck Depression Inventory; STAI = State-Trait Anxiety Inventory; CAR = cortisol awakening response; AUCg = area under the curve with respect to the ground; AUCi = area under the curve with respect to the increase.

The expert consensus guidelines on the CAR by Stalder et al. (2016) indicate that delay between awakening and initiation of sampling is common in CAR studies. In our study, we observed a slight delay in the CAR sampling, with an average delay of 1.83 minutes (SD = 4.75), which is notably shorter than the average delay (7.1 - 24.8 mins) reported by Stalder et al. (2016). Moreover, no significant differences in sampling time delays were found between females and males (p = 0.78), nor between responders and non-responders (p = 0.14).

### 3.2. Associations between chronic social stress, CAR and sex with GMV of ROIs

#### Hippocampus

For both the left and right hippocampi, the models including values from the subscale social tension were significant (left: R^2^ = 0.515, p < 0.001; right: R^2^ = 0.542, p < 0.001; see Table S1 in the supplementary material) with each showing a significant interaction of social tension × sex (left: β = 0.294, p = 0.02; right: β = 0.260, p = 0.035; see Figure 1A-B). To disentangle the interactions, post-hoc partial correlation analyses revealed a significant negative correlation between social tension and hippocampus GMV in males (left: r = −0.375, p = 0.016; right: r = −0.383, p = 0.013), whereas no significant correlations were observed in females (left: r = 0.039, p = 0.814; right: r = 0.014, p = 0.93). Furthermore, Fisher’s Z-test showed that the differences between the correlations in females and males were significant (left: Z = 1.899, p = 0.029; right: Z = 1.831, p = 0.034).

Additionally, for both the left and right hippocampi, the interaction of social tension × CAR was significant (left: β = −0.265, p = 0.034; right: β = −0.237, p = 0.0499; Figure 1C-D). Post-hoc partial correlation analyses showed a negative correlation between social tension and hippocampus GMV for non-responders (left: r = −0.406, p = 0.014; right: r = −0.415, p = 0.012), while the correlation was not significant in responders (left: r = −0.052, p = 0.740; right: r = −0.076, p = 0.628). Fisher’s Z-test showed a significant difference between the correlation coefficients for responders and non-responders of the left hippocampus (Z = −2.11, p = 0.017), and a trend of the right hippocampus (Z = −1.597, p = 0.055).

No other predictor variables or their interactions were significant for hippocampus GMV (see Table S1-S4 for statistical parameters on all regression models).

**Figure 1.**
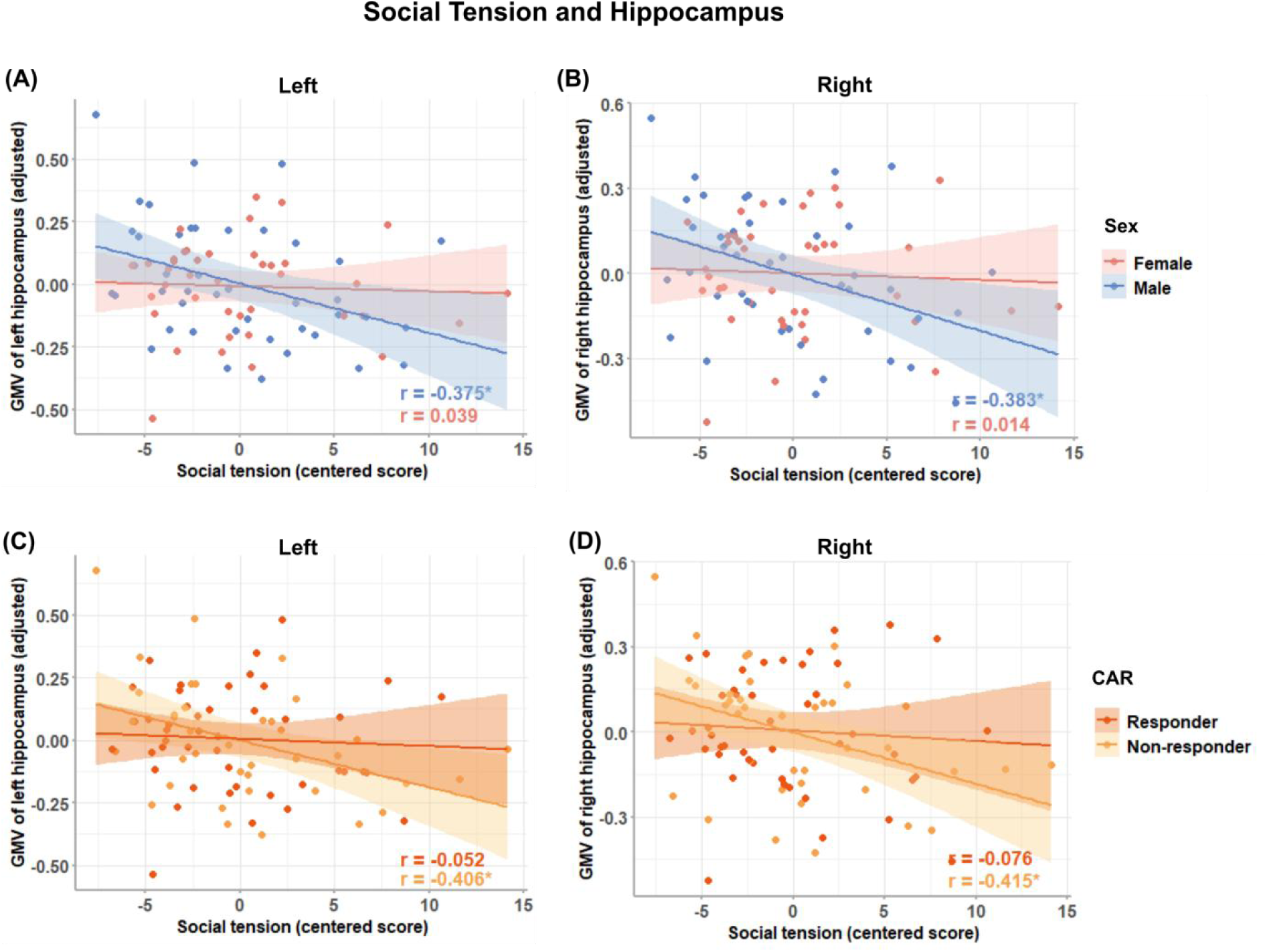
Disentangling the significant interactions of social tension x sex (A-B) and social tension x cortisol awakening response (CAR) (C-D). Males show a significant negative correlation between the amount of socially tense interactions and the hippocampus volume, whereas females show a positive association (left hippocampus: A, right hippocampus: B). Non-responders show a significant negative correlation between socially tense interactions and the hippocampus volume, whereas responders show a positive association (left hippocampus: C; right hippocampus: D).

#### Amygdala

For both the left and right amygdalae, the models including data from the subscale social overload were significant (left: R^2^ = 0.582, p < 0.001; right: R^2^ = 0.609, p < 0.001; see Table S4) with each showing a significant interaction between social overload × CAR (left: β = −0.252, p = 0.032; right: β = −0.279, p = 0.015; Figure 2A-B). Post-hoc partial correlations were not significant for responders (left: r = 0.191, p = 0.220; right: r = 0.192, p = 0.218) and non-responders (left: r = −0.225, p = 0.113; right: r = −0.275, p = 0.105), but Fisher’s Z-tests indicated significant differences between responders and non-responders (left: Z = 1.984, p = 0.024; right: Z = 2.083, p = 0.019) with a positive association in responders and a negative in non-responders (Figure 2A-B). No other predictor or interactions showed a significant association with amygdala GMV (see Table S1-S4 for statistical parameters for all regression models).

**Figure 2.**
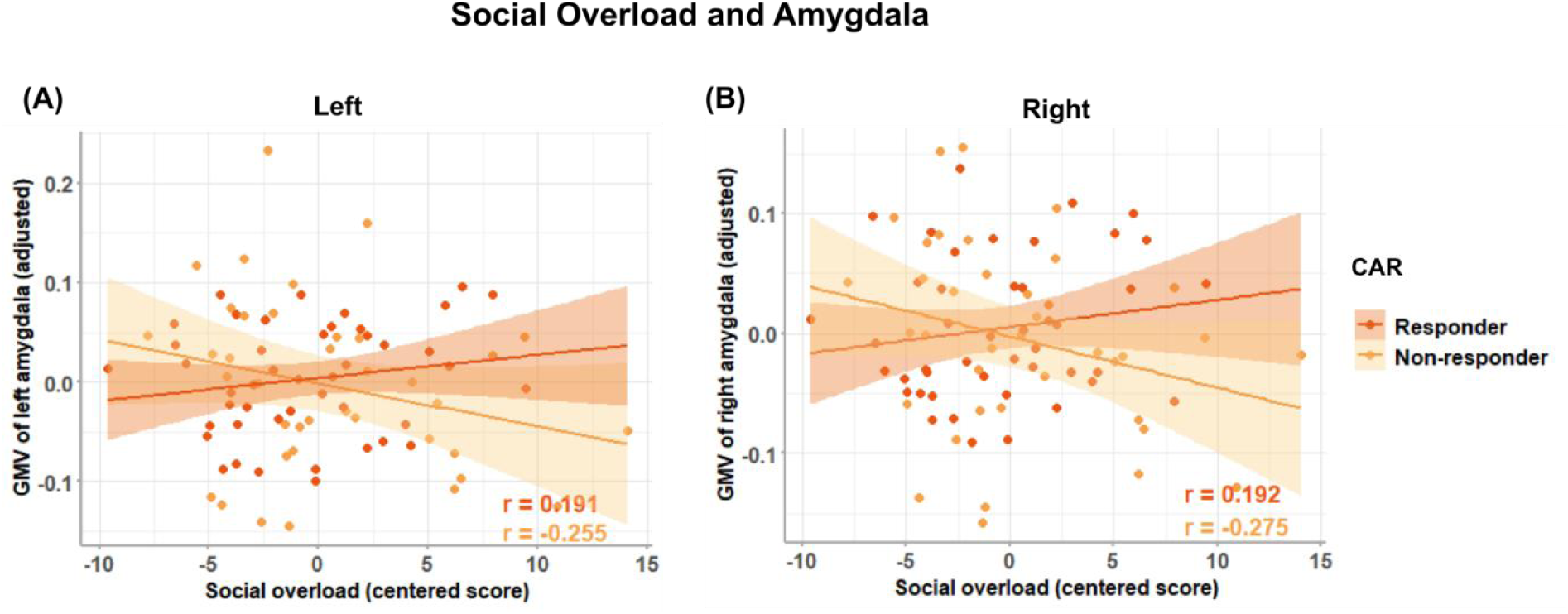
Associations between social overload and GMV of the left (A) and right (B) hippocampus in responders and non-responders. CAR = cortisol awakening response.

## 4. Discussion

The current study examined the impact of different facets of chronic social stress, CAR and sex on hippocampus and amygdala volume in healthy young individuals. Our results show that higher amounts of self-reported socially tense, conflictual interactions are linked to reduced bilateral hippocampus GMV in males, whereas this pattern differs in females. Furthermore, CAR non-responders who report higher amounts of socially tense, conflictual interactions (social tension), and who are overwhelmed by the number of social responsibilities (social overload) show reduced hippocampus and amygdala GMV, whereas CAR responders show the opposite pattern.

### 4.1. The relationship between sex and social stress

Females and males differ in the way social interactions are associated with hippocampus volume, with males who reported higher socially tense interactions showing decreased hippocampus volume. Previous studies with mixed samples of females and males, have consistently shown that high levels of perceived life and social stress are associated with reduced hippocampus volume (e.g., Hanson et al., 2015; Lindgren et al., 2016; Piccolo and Noble, 2018; Zimmerman et al., 2016) and a history of emotional abuse is associated with reduced hippocampus volume in males but not in females (Samplin et al., 2013). This sex-specific pattern aligns well with results reported in animal studies: In male rats, chronic stress was also found to cause dendritic atrophy of hippocampus pyramidal neurons, whereas this was not seen in females (Galea et al., 1997; McLaughlin et al., 2009). In general, prolonged exposure to stress prevents relaxation and maintaining homeostasis, which can subsequently alter brain structure by changing neuronal morphology and suppressing neurogenesis (Kim et al., 2015; Nicolaides et al., 2015). Our results taken together with these reports in the literature point to sex-specific neuro-morphological changes.

One line of argumentation takes steroid hormones such as estradiol and progesterone and their neuroprotective effects into account. Estradiol can enhance dendritic complexity and synaptic spine density in the hippocampus, thereby improving brain volume recovery from injury (Gould et al., 1990; Hoffman et al., 2006; Lee and McEwen, 2001). Progesterone further augments the effect of estradiol (Gould et al., 1990). Both estradiol and progesterone levels are associated with stress reactions in females although not only in a protective way (Aleknaviciute et al., 2017; Bencker et al., 2025; Kirschbaum et al., 1999). Thus, protective effects of steroid hormones against the dendritic retraction in the hippocampus caused by chronic stress (McLaughlin et al., 2010, 2009), might explain the lack of hippocampus volume reduction in socially stressed females.

Additionally, all females in this study were using OC. While controlling for the potential effects of fluctuations in estradiol and progesterone, it is worth mentioning that most OC contain synthetic estradiol such as ethinylestradiol and/or synthetic progesterone such as levonorgestrel. Ethinylestradiol, similar to natural ovarian hormones, induces the synthesis of corticosteroid-binding globulin (CBG) (Moore et al., 1978; Wiegratz et al., 2003), leading to elevated CBG levels during OC use. Since CBG binds to circulating cortisol, the amount of free cortisol available to bind to glucocorticoid receptors in the hippocampus is reduced (Galea et al., 1997; Mataradze et al., 1992), potentially mitigating the negative effects of unbound cortisol on hippocampus volume. Furthermore, OC use has been shown to influence brain structures. Previous studies consistently report a positive association between OC use and total GM volume, as well as an increase in volumes in regions such as the hippocampus, parahippocampus gyrus, and fusiform gyrus, which is also related to the duration and type of OC use (Brouillard et al., 2024; Pletzer et al., 2019, 2015).

Overall, biological protective mechanisms in healthy females appear to prevent negative effects of social stress on hippocampus morphology. This seems to contrast with previous arguments that females are more susceptible to social stress, evidenced by higher cortisol responses to social rejection (Stroud et al., 2002), and that they are more likely to be diagnosed with some stress-related mental disorders (Li and Graham, 2017; Mauvais-Jarvis et al., 2020). Our results revealed that they may show less pronounced volume reductions in the hippocampus and amygdala following chronic stress, while males appear to be more susceptible to stress-related cerebral changes. It is additionally noteworthy that psychosocial stress improves social processing (social cognition, empathy) in males but seem to show no or an opposite effect in females (Nitschke et al., 2022; Smeets et al., 2009). Thus, it might be that social stress such as conflicts also leads to increased empathic stress in males, which is then further affecting cerebral changes in the hippocampus. Our results on social tension highlight the importance of considering sex-specific and neuroendocrine factors in stress-induced brain remodelling.

### 4.2. The association between CAR and social stress

Our data further revealed an interaction of chronic social stress and CAR on brain volume. We observed a significant negative association between social tension and bilateral hippocampus GMV in CAR non-responders, but not in responders. This suggests that individuals who are lacking the typical CAR, i.e., non-responders, might be more vulnerable to the negative impact of socially tense interactions on hippocampus volume. An absent cortisol response to awakening reflects altered neuroendocrine system and impaired HPA axis regulation, which is crucial for maintaining homeostasis as part of the neurobiological circadian recovery system (Stalder et al., 2016). This maladaptive stress response may amplify the negative effects of socially tense interactions on brain structure. In contrast, a functional CAR indicates a neuroprotective, adaptive HPA axis, enabling the individual to effectively adapt to external social stressors and thereby probably mitigate the adverse effects of chronic stress on hippocampus volume.

This pattern is consistent with previous findings that hippocampus lesions and atrophy are associated with attenuated CAR (Buchanan et al., 2004; Pruessner et al., 2007). A healthy cortisol circadian rhythm to awakening – indicated by CAR – may promote adaptive regulation of the HPA axis in response to chronic socially stressful situations. Thus, responders effectively manage their circadian HPA axis activity ensuring restoration and recovery, enabling them to better cope with external social stressors and thereby reducing the negative impact of social tension on the hippocampus. Non-responders, however, seem to lack this protective mechanism, show dysfunctions in the flexibility to adapt to external social stressors and thereby keeping the body’s homeostasis. This might further lead to diminished hippocampus volumes when being chronically exposed to social stressors. Nevertheless, our data so far is only correlational, a causal relationship between the variables needs to be investigated using longitudinal or interventional designs.

The amygdala, a key structure involved in regulating emotions, anxiety, and fear responses (Ressler, 2010; Šimić et al., 2021), is particularly sensitive to psychosocial stress (Roberts et al., 2022) and its functional network is associated with basal cortisol levels (Kogler et al., 2016). Our data indicates that amygdala GMV shows distinct patterns in relation to social overload, differing in CAR responders vs. non-responders. Non-responders compared to responders showed a negative association, with more social overload being associated with reduced amygdala volumes. The association between social overload and amygdala structure was emphasized by a previous study on social media use: Excessive social media use is associated with a reduced bilateral amygdala GMV (He et al., 2017). Our study now additionally indicates that the association between social stress and amygdala volume further depends on the adaptive neurobiological regulation ability of the individual. A healthy circadian neurobiological recovery system even seems to protect from negative impacts of social overload on amygdala volume. This interpretation is further supported by results showing a negative correlation between the left amygdala volume and CAR in patients diagnosed with depression (Roddy et al., 2021). Interestingly, patients with damaged amygdala volume seem to have a smaller social network size (Beadle et al., 2022). Furthermore, no significant interactions with CAR, sex and hippocampus or amygdala volume were found for the dimensions of chronic social isolation and lack of social recognition. The lack of associations may be related to their characteristics, primarily reflecting prolonged social neglect compared to acute conflicts or perceived threats that strongly activate physiological stress responses. Therefore, our findings suggest that social tension and social overload, rather than social neglect, are more prominent psychosocial stressors which in association with the CAR affect hippocampus and amygdala volume.

The dysregulation of the HPA axis is commonly reported in mental disorders (Cherian et al., 2019; Kogler et al., 2025; Wang et al., 2024). In terms of the CAR, previous studies indicate that depression severity is associated with blunted CAR (Adams et al., 2020), while a higher waking cortisol level is related to an increased risk of depression (Adam et al., 2010). Blunted CAR has also been observed in patients with post-traumatic stress disorder (Chida and Steptoe, 2009) and psychosis (Berger et al., 2016). Furthermore, Weber et al. (2023) found that patients with functional neurological disorder, a neuropsychiatric condition characterized by a variety of disabling neurological symptoms, exhibit a blunted CAR, and reduced hippocampus and amygdala volume, which aligns with our findings. Thus, a blunted CAR, along with social stress or emotional neglect, appears to be associated with mental health issues and structural brain changes. Our findings now indicate that even in healthy individuals who do not show significant cortisol increases upon awakening, the hippocampus and amygdala are reshaped by chronic social stress. Therefore, the current study emphasizes the importance of focusing on the adaptability of the stress response in healthy individuals, as indicated by the CAR as an HPA axis function. This further may help preventing negative effects on the neural level and the development of stress-induced mental disorders.

### 4.3. Future directions

Endogenous and exogenous sex hormones that are varying throughout the menstrual cycle, with hormonal contraceptive intake, and during the menopausal transition impact CAR (Høgsted et al., 2021; Stalder et al., 2016). The current study included only females using oral contraceptives which may result in a lower CAR (Høgsted et al., 2021). The impact of endogenous and exogenous sex steroids in females and males at different hormonal stages on CAR and its association with brain structure needs to be addressed in future research (Lee et al., 2024). Second, due to limited resources within the current project, the exact sampling time of CAR relied on self-reports only. CAR samples were collected only on one day, which may not accurately reflect the usual CAR patterns. To improve the validity of collected samples, additional methods to verify sampling and awakening time would be desirable (Richer et al., 2023). Third, while this study focused on structural changes in the hippocampus and amygdala, future research should explore how the network of these brain regions and other regions involved in stress reaction and regulation are affected. This would provide a more comprehensive understanding of stress-related neurobiological mechanisms. Additionally, the different associations between chronic stress and brain structure observed in responders and non-responders in this study need to be further validated in patients with stress-related mental disorders. To identify causal relationships, longitudinal and interventional studies would provide further insights into how sex, chronic social stress and CAR impact brain structure over time.

### 4.4. Conclusion

The present study examined the association between effects of chronic social stress, CAR and sex with GMV in the hippocampus and the amygdala, highlighting the complex interplay of stress-related factors in these brain regions. Our findings demonstrate that chronic social tension is related to hippocampus volume differently in females and males, with a significant negative association between social tension levels and hippocampus GMV only in males. In females, higher levels of ovarian hormones might have a neuroprotective effect and potentially counteract the harmful effects of chronic social stress. In males, increased empathic stress might induce more pronounced anatomical changes. Additionally, we observed that chronic social tension and overload are related to amygdala and hippocampus volume differently in individuals who show a typical cortisol reaction to awakening and those who do not. A significant negative association between stress levels and GMV of amygdala and hippocampus was seen only in non-responders. An inadequate cortisol response upon awakening may reflect underlying dysfunctions in circadian recovery of the HPA axis, potentially increasing the impact of prolonged stress on brain structure. While a functional CAR indicates an adaptive reaction of the body to reallocate and restore the body’s homeostasis to internal and external influencing factors, a missing CAR, as seen in non-responders, indicates a dysfunction in this neurobiological circadian recovery system.

Overall, chronic social stress levels such as conflicts, socially tense and overwhelming interactions are negatively associated with the morphology of the hippocampus and amygdala, particularly among males and CAR non-responders. Understanding the sex-specific relationship between stress and brain structure, as well as the association with adaptive function of the HPA axis, is essential for developing effective stress management and intervention strategies. These insights could help mitigate neural structural changes in individuals who are more vulnerable to social stress and could potentially prevent further development of stress-associated mental disorders.

