## Supplement for "Social stress, cortisol awakening response and sex: Impact on hippocampus and amygdala volume"

**Table S1**. Statistical parameters for the models on social tension, cortisol awakening response (CAR) and sex and grey matter volume (GMV) of the left and right hippocampus and amygdala.

| | **GMV of left hippocampus** | | **GMV of right hippocampus** | | **GMV of left amygdala** | | **GMV of right amygdala** | |
|---|---|---|---|---|---|---|---|---|
| | **β** | **t** | **β** | **t** | **β** | **t** | **β** | **t** |
| Age | 0.05 | 0.58 | 0.01 | 0.15 | 0.10 | 1.31 | 0.07 | 0.85 |
| TIV | 0.54 | **4.55***** | 0.63 | **5.44***** | 0.56 | **5.09***** | 0.59 | **5.44***** |
| Sex | -0.12 | -0.84 | -0.05 | -0.31 | -0.18 | -1.30 | -0.17 | -1.24 |
| CAR | -0.05 | -0.44 | -0.03 | -0.27 | -0.02 | -0.17 | -0.06 | -0.53 |
| Social tension | -0.20 | -1.50 | -0.20 | -1.50 | -0.16 | -1.32 | -0.09 | -0.73 |
| Social tension × sex | 0.29 | **2.36*** | 0.26 | **2.14*** | 0.18 | 1.59 | 0.18 | 1.60 |
| Social tension × CAR | -0.27 | **-2.16*** | -0.24 | **-1.99*** | -0.14 | -1.22 | -0.13 | -1.13 |
| Sex × CAR | 0.04 | 0.31 | 0.01 | 0.10 | -0.04 | -0.32 | -0.01 | -0.05 |
| | $R^2 = 0.515$, $p < 0.001$ | | $R^2 = 0.552$, $p < 0.001$ | | $R^2 = 0.589$, $p < 0.001$ | | $R^2 = 0.591$, $p < 0.001$ | |

Note: TIV = total intracranial volume. * indicates predictors significant at p < 0.05; *** indicates predictors significant at p < 0.001.

**Table S2.** Statistical parameters for the models on social overload, CAR and sex and GMV of the left and right hippocampus and amygdala.

| | **GMV of left hippocampus** | | **GMV of right hippocampus** | | **GMV of left amygdala** | | **GMV of right amygdala** | |
|---|---|---|---|---|---|---|---|---|
| | **β** | **t** | **β** | **t** | **β** | **t** | **β** | **t** |
| Age | 0.06 | 0.67 | 0.02 | 0.24 | 0.11 | 1.38 | 0.09 | 1.14 |
| TIV | 0.63 | **5.06***** | 0.71 | **5.87***** | 0.62 | **5,70***** | 0.66 | **6.31***** |
| Sex | -0.03 | -0.22 | 0.04 | 0.25 | -0.10 | -0.77 | -0.10 | -0.73 |

| | | | | | | | | |
|---|---|---|---|---|---|---|---|---|
| CAR | 0.01 | 0.05 | 0.02 | 0.19 | 0.03 | 0.26 | -0.01 | -0.09 |
| Social overload | 0.06 | 0.39 | 0.04 | 0.31 | 0.12 | 0.96 | 0.10 | 0.78 |
| Social overload × sex | 0.08 | 0.69 | 0.08 | 0.70 | 0.03 | 0.27 | 0.09 | 0.92 |
| Social overload × CAR | -0.23 | -1.76 | -0.21 | -1.65 | -0.25 | **-2.19*** | -0.28 | **-2.50*** |
| Sex × CAR | -0.02 | -0.22 | -0.05 | -0.35 | -0.09 | -0.71 | -0.05 | -0.39 |
| | $R^2 = 0.456$, $p < 0.001$ | | $R^2 = 0.049$, $p < 0.001$ | | $R^2 = 0.582$, $p < 0.001$ | | $R^2 = 0.609$, $p < 0.001$ | |

Note: * indicates predictors significant at $p < 0.05$; *** indicates predictors significant at $p < 0.001$.

**Table S3**. Statistical parameters for the models on lack of social recognition, CAR and sex and GMV of the left and right hippocampus and amygdala.

| | **GMV of left hippocampus** | | **GMV of right hippocampus** | | **GMV of left amygdala** | | **GMV of right amygdala** | |
|---|---|---|---|---|---|---|---|---|
| | **β** | **t** | **β** | **t** | **β** | **t** | **β** | **t** |
| Age | 0.03 | 0.38 | -0.01 | -0.07 | 0.08 | 1.07 | 0.06 | 0.79 |
| TIV | 0.60 | **4.84***** | 0.69 | **5.72***** | 0.61 | **5.64***** | 0.65 | **6.03***** |
| Sex | -0.07 | -0.42 | 0.01 | 0.07 | -0.13 | -0.93 | -0.13 | -0.97 |
| CAR | 0.02 | 0.12 | 0.03 | 0.25 | 0.05 | 0.45 | -0.01 | -0.12 |
| Lack of social recognition | -0.08 | -0.56 | -0.03 | -0.20 | -0.03 | -0.24 | 0.02 | 0.19 |
| Lack of social recognition × sex | 0.15 | 1.26 | 0.11 | 0.95 | 0.10 | 0.99 | 0.10 | 0.92 |
| Lack of social recognition × CAR | -0.13 | -1.05 | -0.15 | -1.28 | -0.20 | -1.87 | -0.18 | -1.67 |

| | | | | | | | | |
|---|---|---|---|---|---|---|---|---|
| Sex × CAR | -0.02 | -0.13 | -0.05 | -0.32 | -0.10 | -0.78 | -0.03 | -0.26 |
| | $R^2$ = 0.451, p < 0.001 | | $R^2$ = 0.486, p < 0.001 | | $R^2$ = 0.587, p < 0.001 | | $R^2$ = 0.592, p < 0.001 | |

Note: *** indicates predictors significant at p < 0.001.

**Table S4**. Statistical parameters for the models on social isolation, CAR and sex and GMV of the left and right hippocampus and amygdala.

| | **GMV of left hippocampus** | | **GMV of right hippocampus** | | **GMV of left amygdala** | | **GMV of right amygdala** | |
|---|---|---|---|---|---|---|---|---|
| | **β** | **t** | **β** | **t** | **β** | **t** | **β** | **t** |
| Age | 0.05 | 0.57 | 0.01 | 0.09 | 0.10 | 1.24 | 0.08 | 1.00 |
| TIV | 0.60 | **4.78*** | 0.68 | **5.58*** | 0.60 | **5.44*** | 0.63 | **5.85*** |
| Sex | -0.08 | -0.48 | 0.00 | 0.00 | -0.13 | -0.97 | -0.13 | -0.99 |
| CAR | -0.04 | -0.34 | -0.01 | -0.12 | -0.01 | -0.12 | -0.05 | -0.50 |
| Social isolation | -0.01 | -0.06 | -0.01 | -0.05 | 0.08 | 0.64 | -0.03 | -0.22 |
| Social isolation × sex | -0.05 | -0.44 | -0.02 | -0.18 | -0.05 | -0.45 | 0.01 | 0.14 |
| Social isolation × CAR | 0.12 | 1.01 | 0.05 | 0.41 | 0.07 | 0.67 | 0.07 | 0.71 |
| Sex × CAR | 0.04 | 0.25 | -0.01 | -0.06 | -0.05 | -0.34 | 0.01 | 0.05 |
| | $R^2$ = 0.437, p < 0.001 | | $R^2$ = 0.467, p < 0.001 | | $R^2$ = 0.564, p < 0.001 | | $R^2$ = 0.574, p < 0.001 | |

Note: *** indicates predictors significant at p < 0.001.
